# The parasitic worm product ES-62 normalises the gut microbiota/bone marrow axis in inflammatory arthritis

**DOI:** 10.1101/459768

**Authors:** James Doonan, Anuradha Tarafdar, Miguel A. Pineda, Felicity E. Lumb, Jenny Crowe, Aneesah M. Khan, Paul A. Hoskisson, Margaret M. Harnett, William Harnett

## Abstract

The human immune system has evolved in the context of our colonisation by bacteria, viruses, fungi and parasitic helminths. Reflecting this, the rapid eradication of pathogens appears to have resulted in reduced microbiome diversity and generation of chronically activated immune systems, presaging the recent rise of allergic, autoimmune and metabolic disorders. Certainly, gastrointestinal helminths can protect against gut and lung mucosa inflammatory conditions by modulating the microbiome and suppressing the chronic inflammation associated with dysbiosis. Here, we employ ES-62, an immunomodulator secreted by tissue-dwelling *Acanthocheilonema viteae* to show that modulation of the gut microbiome does not require live infection with gastrointestinal-based helminths nor is protection restricted to mucosal diseases. Specifically, subcutaneous administration of this defined parasitic worm product affords protection against joint disease in collagen-induced arthritis, a mouse model of rheumatoid arthritis, which is associated with normalisation of gut microbiota and prevention of loss of intestinal barrier integrity.

## 1. Introduction

Parasitic helminths (worms) have evolved the ability to modulate host immune and tissue repair responses in order to promote their survival by limiting the inflammation that would otherwise drive their expulsion and cause pathology^1^. The recent rapid eradication of helminths (and other infectious pathogens) appears to have resulted in over-activated immune systems and this provides a rationale for the increasing prevalence of chronic allergic (e.g. asthma) and autoimmune (e.g. inflammatory bowel disease, type-1 diabetes, multiple sclerosis [MS], systemic lupus erythematosus [SLE] and rheumatoid arthritis [RA]) inflammatory disorders, as well as contributing to the rise in obesity and associated metabolic syndrome co-morbidities including type-2 diabetes and cardiovascular disease^2-6^ in developing and urbanised countries. Although genetic studies of patients have identified variants of genes that are associated with these inflammatory diseases in many cases such as RA, these alone do not appear to be strong risk factors. Rather, integration with signals driven by environmental factors (e.g. smoking or diet) is required to trigger disease. Recognition of the impact of environmental factors has focused interest on the role of the microbiota^5,7,8^ and hence, on how helminths may regulate this in health and disease^3,9,10^. Indeed, emerging data suggest that commensal bacteria and gastrointestinal (GI) helminths appear to reciprocally regulate the composition of the gut microbiome^11,12^. It appears that such crosstalk has evolved to homeostatically maintain immune system function in health and disease^2^ as GI helminths can induce regulatory responses to limit inflammation and promote intestinal barrier integrity while the intestinal bacteria play an essential role in training the immune system by impacting on stem and progenitor cells^13,14^. Consistent with the former, there is increasing evidence from animal models that the protection afforded by GI helminth infection, against inflammatory disorders associated with mucosal tissues like asthma^15^ and inflammatory bowel disease^16,17^ and coeliac disease^18,19^, involves modulation of the gut microbiota.

Nevertheless, gut, lung or oral dysbiosis has also been implicated in the aetiology of a wide range of systemic and organ-specific autoimmune diseases including musculoskeletal pathologies like RA and SLE^20-31^. It is not clear whether the protection afforded by GI helminths against these disorders similarly involves interaction with the microbiome, but infection with helminths like *Heligmosomoides polygyrus* and *Trichuris muris* can result in increases in *Lactobacillaceae* and decreases in *Prevotella* species^32-35^, commensals that are reported to be dysregulated in RA patients^24,31^. In any case, helminth-mediated protection against autoimmune disease is not limited to GI-tract parasites, with particularly striking examples of this involving filarial nematodes with respect to RA^36^ and SLE^37^ being reported in India. However, to date it has been unclear whether tissue-resident or blood-borne parasitic worms can mediate these effects via modulation of the host microbiome and if so, which mechanisms they utilise to achieve this protection.

The ability of helminths to ameliorate chronic inflammatory disorders has often been attributed to their capacity to excrete or secrete molecules (ES) that exert immunoregulation to promote immune-evasion and limit host pathology^2^. Amongst the best characterised ES products is ES-62, a phosphorylcholine (PC)-containing glycoprotein secreted by the rodent filarial nematode *Acanthocheilonema viteae* that we have shown to prevent initiation and progression of pathology in mouse models of certain allergic (asthma, contact dermatitis) and autoimmune (RA, SLE) inflammatory diseases^1,2,38-43^. Collectively, our studies have identified a unifying mechanism of action that allows effective protection irrespective of the inflammatory phenotype: thus, by subverting TLR4 signalling to downregulate aberrant MyD88-responses, ES-62 acts to homeostatically reset the regulatory:effector immune cell balance, primarily to restore levels of IL-10+ regulatory B cells (Bregs) and suppress pathological IL-17-driven inflammation^1,2,38-44^. In experimental models of RA and human disease, perturbation of the microbiota has been shown to disrupt the balance of pathogenic Th17 cells and the counter-regulatory Bregs and Tregs that act to homeostatically resolve inflammation^20,24-26^. Thus, our aim here was to investigate whether the actions of ES-62 reflected an ability to impact on the microbiome. Consistent with this, we now show that whilst joint disease in the collagen-induced arthritis (CIA) mouse model of RA is preceded by disturbance of the gut microbiome with accompanying intestinal inflammation and loss of barrier integrity, ES-62 acts to normalise the microbiome and maintain gut health and function. Further supporting our hypothesis that such local gut pathology and inflammation drives joint disease, prophylactic depletion of the gut microbiota with broad-spectrum antibiotics (ABX) was found to reduce the consequent severity of arthritis in mice undergoing CIA. Such ABX-treatment also reduced the level of protection afforded by ES-62, with these animals exhibiting an intermediate phenotype of disease, comparable to that of the control ABX-treated mice undergoing CIA. These data therefore indicate that a “normalised” microbiome is also required for the full induction of the immunoregulatory actions of ES-62.

## 2. Materials and Methods

### 2.1. Collagen induced Arthritis (CIA)

Male DBA/1 mice were purchased at 6-8 weeks of age (Envigo; Bicester, UK) and then housed and maintained in the Central Research Facility of the University of Glasgow. All experiments were approved by, and conducted in accordance with, the Animal Welfare and Ethical Review Board of the University of Glasgow, UK Home Office Regulations and Licenses PPL P8C60C865, PIL I518666F7, PIL 1675F0C46 and PIL ICEBDB864. CIA was induced using bovine Collagen Type II (CII ‐ 100 μg) emulsified with complete Freud’s adjuvant (MD Biosciences) injected intradermally on day 0. Mice were challenged with 200 μg CII in PBS intraperitoneally on day 21. Animals were treated with PBS or purified endotoxin-free ES-62 (2 μg/injection) subcutaneously on days ‐2, 0 and 21 and joint inflammation and damage (articular score) determined as described previously^39,45,46^. Grip strength was recorded as per manufacturer’s instruction (Ugo basile®, Italy) using a Grip-Strength Meter, which measured the grip strength (peak force and time resistance) of the forelimbs of the mice. The animals were placed over a base plate and gripped a T-shaped grasping bar which was connected to the peak amplifier that automatically detects the animal’s response. Three measurements were taken and the average grip strength was calculated. In order to investigate the impact of gut microbiome perturbation on initiation and progression of inflammatory arthritis, animals were given drinking water containing [or not] a cocktail of antibiotics (500 mg/L Vancomycin, 1g/L Neomycin and 1g/L Metronizadole) to eliminate Gram-positive, Gram negative and anaerobic microorganisms ^25^ 7 days prior to the induction of CIA and thereafter continuously throughout the experiment. Blood was sampled using endotoxin-free needles and syringes and the resulting serum isolated and stored at ‐20^°^C in endotoxin-free Eppendorf tubes. Paws, ileum and colon tissue were fixed in 4% paraformaldehyde; ileum and colon faecal contents were collected in sterile RNAlater (Sigma) and stored at ‐80^°^C.

### 2.2. Flow Cytometry

Spleen and bone marrow (BM) cells were suspended in FACS buffer (2.5% BSA; 0.5 mM EDTA, in PBS) following red blood cell-lysis (eBioscience). BM cells were labelled with a cocktail of PE-labelled antibodies specific for CD3, B220 and Ter119 to exclude analysis of lymphocytes and erythroid cell populations using a dump channel, and monocytes were identified by labelling with antibodies against CD11b (FITC), Ly6C (PerCP Cy5.5) and Ly6G (APC)^43^. Lymphocytes from the spleen were labelled with antibodies specific for CD19 (AF700) and IL-10 (PE or APC). All antibodies were purchased from BioLegend, UK. Fixable viability stain (APC-ef780; ThermoFisher Scientific) was used to select for live cells and for analysis of IL-10^+^ regulatory B cells (Bregs), lymphocytes were stimulated with PMA, ionomycin, Brefeldin A and LPS as described previously^41,44^. Data were acquired using a FACS Canto or BD LSRII flow cytometer and populations were gated as described previously using isotype and fluorescence minus one (FMO) controls using FlowJo, LLC analysis software (Tree Star/ BD)^41,44,43^.

### 2.3. Histology

Ileum, colon and joint (paw) tissues from individual mice in each treatment group were fixed in 4% paraformaldehyde for 24 hours before gut tissues were embedded in OCT and paw joints were decalcified and subsequently paraffin embedded. Paraffin sections (6 μm) and OCT cryosections (9-10 μm) were prepared and standard H&E histological staining was performed on all tissues for identification of morphological changes^39,43,46^. Ileum (villi thickness) and colon (number of lesions) pathology was quantitated by Image J analysis. Joint pathology was scored according to the grading system of 0 for no inflammation, 1 for mild inflammation, pannus formation and bone damage, up to a score of 4 representing a high level of inflammation, pannus infiltration and bone and cartilage destruction, as previously described^47^.

### 2.4. Osteoclast differentiation

OCs were differentiated from BM obtained from the hind limbs of experimental animals as previously described^43^. Briefly, following removal of adherent cells, BM cells were cultured in αMEM medium supplemented with 30 ng/ml M-CSF and 50 ng/ml RANKL (Peprotech, London, UK) and then assessed for OC differentiation by TRAP staining (Leukocyte Acid Phosphatase Kit, Sigma-Aldrich, UK) on day 5. Images were obtained using an EVOS FL Auto Cell Imaging System. TRAP^+^ cells with ≥ 3 nuclei were enumerated and Image J software was used to calculate the average size of multinucleated OCs per field of view (FoV)^43^.

### 2.5. Serum cytokine and antibody ELISAs

Interleukin-6 (IL-6) and IL-10 expression was measured by ELISA according to the manufacturer’s instructions (BD Biosciences, Oxford, UK). For determination of collagen type II (CII)-specific IgG1 and IgG2a antibodies in serum^45^, high binding 96 well ELISA plates were coated with CII (5 μg/ml) overnight at 4^°^C before washing and blocking with BSA/PBS. Serum was diluted 1:100 and then serially diluted three-fold until 1:218700 and incubated with HRP-conjugated goat anti-mouse IgG1 or IgG2a (1:10,000) in 10% FBS/PBS prior to developing with TMB and 2M sulphuric acid and read at an optical density of 450nm.

### 2.6. qRT-PCR

BM cells (10^6^) were lysed in RLT Lysis Buffer, while splenic tissue was lysed in Trizol reagent (Sigma), prior to mRNA extraction using RNeasy Plus Mini kit (Qiagen, Germany) or phenol-chloroform extraction, respectively, according to the manufacturer’s instructions. The High Capacity cDNA Reverse Transcriptase kit (Applied Biosystems, Life Technology, UK) was used to generate cDNA for use with StepOne Plus^TM^ real-time PCR system (Applied Biosystems, UK) and KiCqStart^®^ qPCR Ready Mix (Sigma-Aldrich). Pre-designed KiCqStart^TM^ Primers (Sigma-Aldrich) were purchased to evaluate RANK (*tnfrsf11a*; forward – GAAATAAGGAGTCCTCAGGG, reverse ‐ GAAATAAGGAGTCCTCAGGG), OPG (*tnfrsf11b*; forward – GAAGATCATCCAAGACATTGAC, reverse ‐ TCCTCCATAAACTGAGTAGC), MyD88 (*myd88*; forward – GAAGATCATCCAAGACATTGAC, reverse ‐ TCCTCCATAAACTGAGTAGC) and β-actin (*actb*; forward – GATGTATGAAGGCTTTGGTC, reverse ‐ TGTGCACTTTTATTGGTCTC). Data were normalised to the reference gene β-actin to obtain the ΔCT values that were used to calculate the fold-change from the ΔΔCT following normalisation to biological control group.

### 2.7. Metagenomics

Genomic DNA from the ileum and colon faecal matter was purified using QIAamp DNA Stool Mini Kit (Qiagen, UK) and stored at ‐20^°^C. For metagenomic analysis using the Ion Torrent PGM™ platform, samples from three individual mice per group were pooled and between 10 and 100ng of the pooled DNA was fragmented (NEB Fast DNA Fragmentation & Library Prep Set for Ion Torrent, NEB Inc, UK) and barcoded (IonXpress Barcode Adapters Kit, ThermoFisher Scientific, UK). Barcoded libraries were quantified using Qubit Fluorometer (ThermoFisher Scientific, UK) and bioanalyser (High Sensitivity DNA analysis Kit, Agilent, UK). Up to three barcoded libraries were combined per Ion 316^TM^ Chip Kit v2 following library preparation using the Ion PGM^TM^ Hi-Q^TM^ View OT2 and Ion PGM^TM^ Hi-Q^TM^ View Sequencing Kits (ThermoFisher Scientific, UK). Data were extracted as FASTQ files and analysed using MG-RAST to generate taxonomic data from sequencing reads^48^. The number of reads per phylum, class, order, family, genera or species of interest were expressed as a composition of all bacteria present to normalise for variation between sequencing runs. Sequencing runs can be accessed using MG-RAST IDs; mgm4777616.3, 4777615.3, 4777614.3, 4777613.3, 4777481.3, 4777480.3, 4777479.3, 4777478.3, 4767994.3, 4767993.3, 4767992.3, 4767991.3, 4767990.3, 4767989.3, 4767988.3, 4767987.3, 4767986.3, 4738191.3, 4738190.3, 4738025.3, 4737887.3, 4737053.3 and 4737052.3. qPCR was used to validate changes in bacterial populations using primers specific for Bacteriodetes (forward ‐ GTTTAATTCGATGATACGCGAG, reverse ‐ TTAAGCCGACACCTCACGG) Firmicutes (forward ‐ GGAGCATGTGGTTTAATTCGAAGCA, reverse ‐ AGCTGACGACAACCATGCAC) and *Butyrivibrio* (forward – GCGAAGAAGTATTTCGGTAT, reverse ‐ CCAACACCTAGTATTCATC) and were normalised to the total levels of bacteria using pan-bacterial primers (forward ‐ CGGTGAATACGTTCCCGG, reverse ‐ TACGGCTACCTTGTTACGACTT).

### 2.8. Statistics

All data were analysed using GraphPad Prism 6 software using One or Two-Way ANOVA with Fishers LSD post-tests for parametric data or Kruskal-Wallis test and Dunn’s post-test for non-parametric data. Unsupervised hierarchical clustering with Euclidean distance was performed on the colon samples using the heatmap.2 function of the gplots package in R. Supervised heatmaps were generated using GraphPad Prism 7 software. Indicators of significance include * = p < 0.05, ** = p < 0.01 and *** = p < 0.001.

## 3. Results

### 3.1 ES-62 protection against CIA is associated with normalisation of the gut microbiome

As reported previously, ES-62 ameliorates CIA in terms of articular score (Fig. 1a) and reflecting this protection against joint morbidity, we now show that it prevents the resulting low grip strength found in CIA-mice (Fig. 1b). Indeed, treatment with ES-62 maintains grip strength in CIA-mice at a similar level to that found in healthy, Naive (not subjected to CIA) DBA/1 mice (Fig. 1b). Commensal bacteria have increasingly been proposed to play a role in RA pathogenesis (reviewed in^49-51^) and the decline in grip strength during ageing has been associated with changes in the gut microbiome^52^. Interestingly, in addition to being an indicator of frailty, grip strength has been shown to be a predictor of a wide range of adverse health outcomes^53^, e.g., cardiovascular disease, which RA patients are at increased risk of developing^54^ and that are impacted by the microbiome^49,55^. Thus, to address whether the protection afforded by ES-62 is associated modulation of the gut microbiota, a metagenomic approach was used to profile the bacterial populations present in the intestines of CIA-mice, exposed or not to ES-62. Initiation of RA (and CIA) pathogenesis is associated with disruption of the balance of effector:regulatory immune system cells and so we characterised the bacterial changes pertaining during established arthritis in the ileum and colon, intestinal sites where the microbiome and the metabolic microenvironment play key roles in shaping Th17 (ileum-associated) and regulatory (colon-associated) immune responses^25,56,57^.

**Fig. 1.**
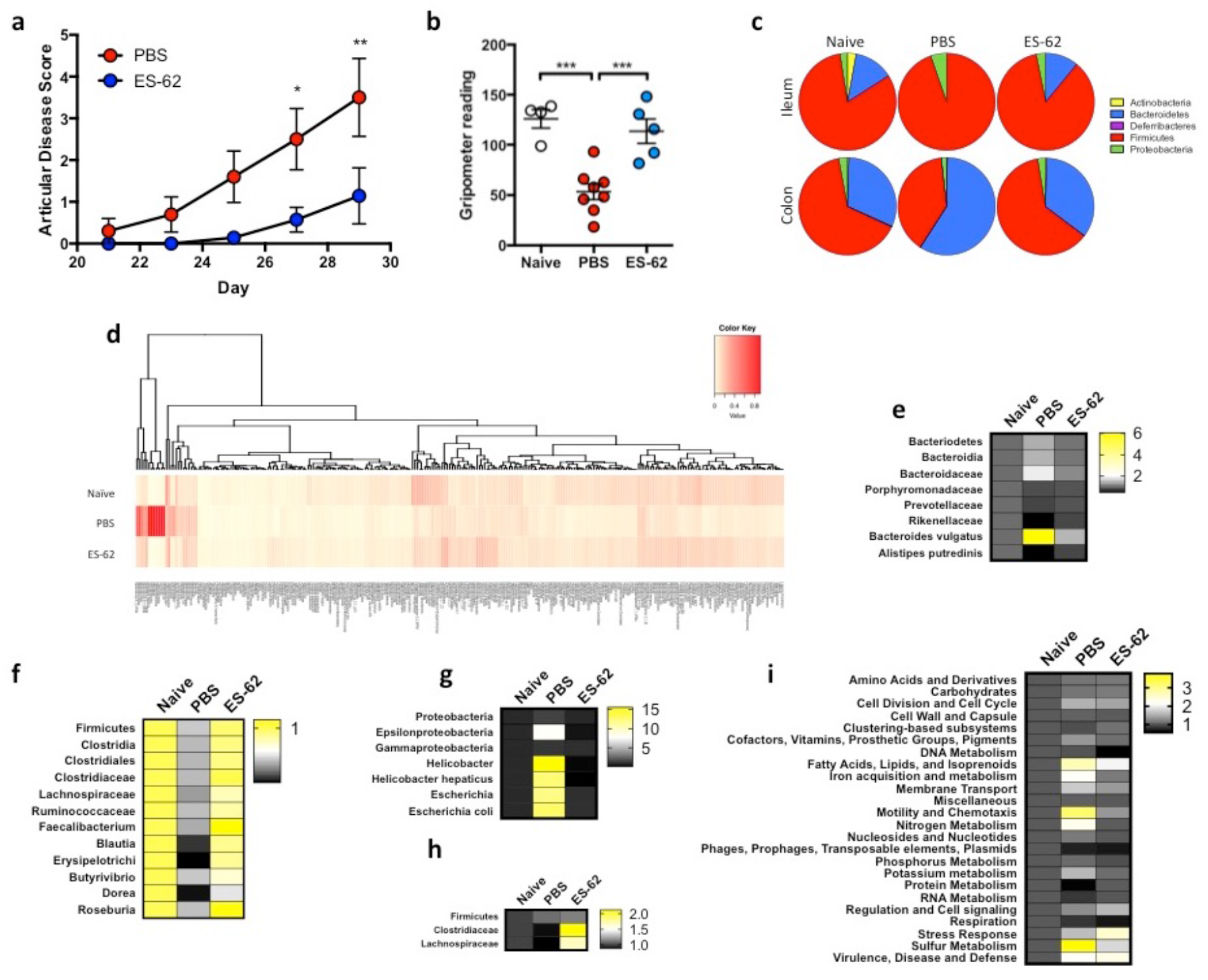
ES-62 “normalises” the microbiome during CIA to a naïve phenotype. (**a**) Articular scores expressed as means ± SEM of PBS (n = 10) or ES-62 (n = 7) ‐treated CIA animals, with data pooled from 3 experiments. (**b**) Grip strength expressed as an average of three measurements per mouse measured prior to cull as an indicator of forelimb strength and joint morbidity (Naïve; n = 4, PBS; n = 8 and ES-62; n = 6). (**c**) The composition of bacterial phyla present in the ileum and colon of Naïve, PBS and ES-62-treated CIA animals presenting proportion values as pie charts from a single representative experiment using pooled samples from 3 mice in each condition. (**d**) Heatmap analysis of all bacteria present in the colon of Naïve, PBS and ES-62-treated CIA animals (n=3/group) from a representative model is shown. (**e - i**) Statistically significant changes between the PBS-CIA and ES-62-CIA groups in Bacteriodetes (**e**; Bacteriodaceae; PBS versus Naïve or ES-62, p < 0.05), Firmicutes (**f;** Clostridiales; PBS versus Naïve, p < 0.01 and PBS versus ES-62, p < 0.05) and Proteobacteria (**g**; Epsilonbacteria; PBS versus Naïve or ES-62, p < 0.05) in the colon and Firmicutes in the ileum (**h**; Clostridiaceae; PBS versus ES-62, p < 0.05) as well as functional metagenomics of the colon (**i**; Phages; Naive versus PBS, p < 0.01, Protein metabolism; PBS versus Naïve or ES-62, p < 0.01) are presented as heatmaps with changes in bacterial populations in PBS‐ and ES-62-treated CIA animals normalised to Naïve controls. Three mice per group were pooled for metagenomic analysis and the mean data from 3 independent experiments are presented. Statistical significance was determined using Two-way (**a, e - i**) and one-way ANOVA (**b**) with LSD Fishers multiple comparisons and for levels of significance where indicated by asterisks, * = p < 0.05, ** = p < 0.01 and *** = p < 0.001.

An overview of the microbiota of the ileum and colon at the phylum level shows substantial changes between healthy Naive DBA/1 mice and those with established arthritic disease (PBS; Fig. 1c). Firmicutes and Bacteriodetes are the predominant phyla in all treatment groups but with respect to CIA-mice, whereas they exhibit outgrowths of Firmicutes and Proteobacteria in the ileum, by contrast they exhibit decreased levels of Firmicutes with a compensatory outgrowth of Bacteriodetes in the colon (Fig. 1c). ES-62 essentially helps to maintain the healthy diversity of the microbiota observed in Naive mice, which was reduced in the CIA-mice (Fig. 1c). Deeper analysis illustrates the differential diversity signatures exhibited by healthy and arthritic mice, as well as the impact of ES-62 treatment (Fig. 1d): drilling down on the modulation of the Gram −ve Bacteroidetes phylum reveals some key differential signatures throughout each of the predominant *Bacteroides*, *Porphyromonas* and *Prevotella* genera between the colon contents of Naive and CIA mice, and identifying those normalised by exposure to ES-62 (Fig. 1e). Strikingly, at the species level, *B. vulgatus* was found to be significantly increased in the colon of CIA relative to Naive or ES-62-CIA mice (Fig. 1e). By contrast abundance of members of the *Rikenellaceae* family and specifically, *Alistipes putredinis*, were reduced in the colon of CIA mice when compared to Naive and ES-62-treated CIA mice (Fig. 1e). Similarly, in spite of the fact that differential profiles amongst the treatment groups were observed throughout major genera (e.g. *Bacillus*, *Staphyloccus*, *Streptococcus*, *Enterococcus* and *Clostridium*) of Gram +ve Firmicutes, the most dramatic changes observed with established CIA occurred within the Clostridiales order. In particular, decreases in the *Clostridiaceae* (Fig. 1f) and the *Lachnospiraceae* (Fig. 1f) families were noted with ES-62 promoting maintenance of the *Ruminococcus*, *Faecalibacterium*, *Blauti* and *Erysipelotrichaceae* genera and butyrate-producing *Dorea* and *Roseburia* species, the latter of which have been implicated in gut health and inflammation homeostasis^58^. In terms of the Proteobacteria, CIA was associated with outgrowth of members of the *Helicobacter*, specifically *Helicobacter hepaticus* (Epsilonproteobacteria) and *Escherichia*, particularly *Escherichia coli* (Gammaproteobacteria) genera (Fig. 1g) and again this was normalised by ES-62.

Perturbation of the microbiome was also observed in the ileum of CIA mice and again, exposure to ES-62 acted to normalise this towards the healthy community (Fig. 1c). In terms of protective signatures, despite the relative paucity of bacteria in the ileum relative to the colon, ES-62 clearly promoted growth of the *Clostridiales*, again particularly *Clostridaceae* and *Lachnospiraceae* species but in this case generally promoting their outgrowth beyond the levels found in healthy mice (Fig. 1h). This presumably reflects that whilst treatment of naive, healthy DBA/1 mice with ES-62 for the duration of the CIA model had little effect on the colon microbiota, it promoted expansion of *Clostridiales* species in the ileum (Fig. S1a and b), perhaps explaining the increase in species diversity observed in the ileum (Fig. S1c) but not the colon (Fig. S1d) of CIA-mice treated with ES-62. Additionally, reflecting the ability of CIA to perturb, and ES-62 to normalise, the gut bacteria, functional metagenomic analysis also showed that ES-62 generally acted to normalise the metabolic capacity of the colonic microbiome (Fig. 1i).

### 3.2. Perturbation of the gut microbiome with broad-spectrum antibiotics both ameliorates CIA *and* impacts on the protection afforded by ES-62

To address whether the perturbation of the microbiome observed in CIA plays a role in the initiation and progression of inflammatory arthritis, we investigated the effect of continuous exposure of mice to a cocktail of broad-spectrum antibiotics (ABX; 500 mg/L Vancomycin, 1g/L Neomycin and 1g/L Metronizadole) administered in their drinking water from one week prior to initiation of CIA. Such ABX treatment had no obvious effect on the overall health of the animals as, after a characteristic initial dip, there was no significant difference in body weight amongst the CIA groups at cull (Fig. S2a). Nevertheless, this regimen essentially eliminated the bacterial microbiota of all animals irrespective of treatment group (Fig. S2b), whilst metagenomic analysis showed the residual gut community to be almost entirely comprised of proteobacteria (Fig. S2c and S2d). As predicted from previous studies in a variety of inflammatory arthritis models^25^,^59-61^, ABX treatment reduced the incidence (PBS, 65.2%; PBS-ABX, 36.3%, as measured by articular score ≥ 1) and severity of joint pathology in CIA-mice, both in terms of articular score (Fig. 2a) and histopathology (Fig. 2b and c). In addition, the protection afforded by ES-62 was reduced in ABX-treated animals such that the articular score of such mice was not significantly different from PBS-CIA mice receiving ABX or not (Fig 2a). Thus, prophylactic administration of ABX resulted in an intermediate phenotype of CIA irrespective of whether the mice were treated with PBS or ES-62.

**Fig. 2.**
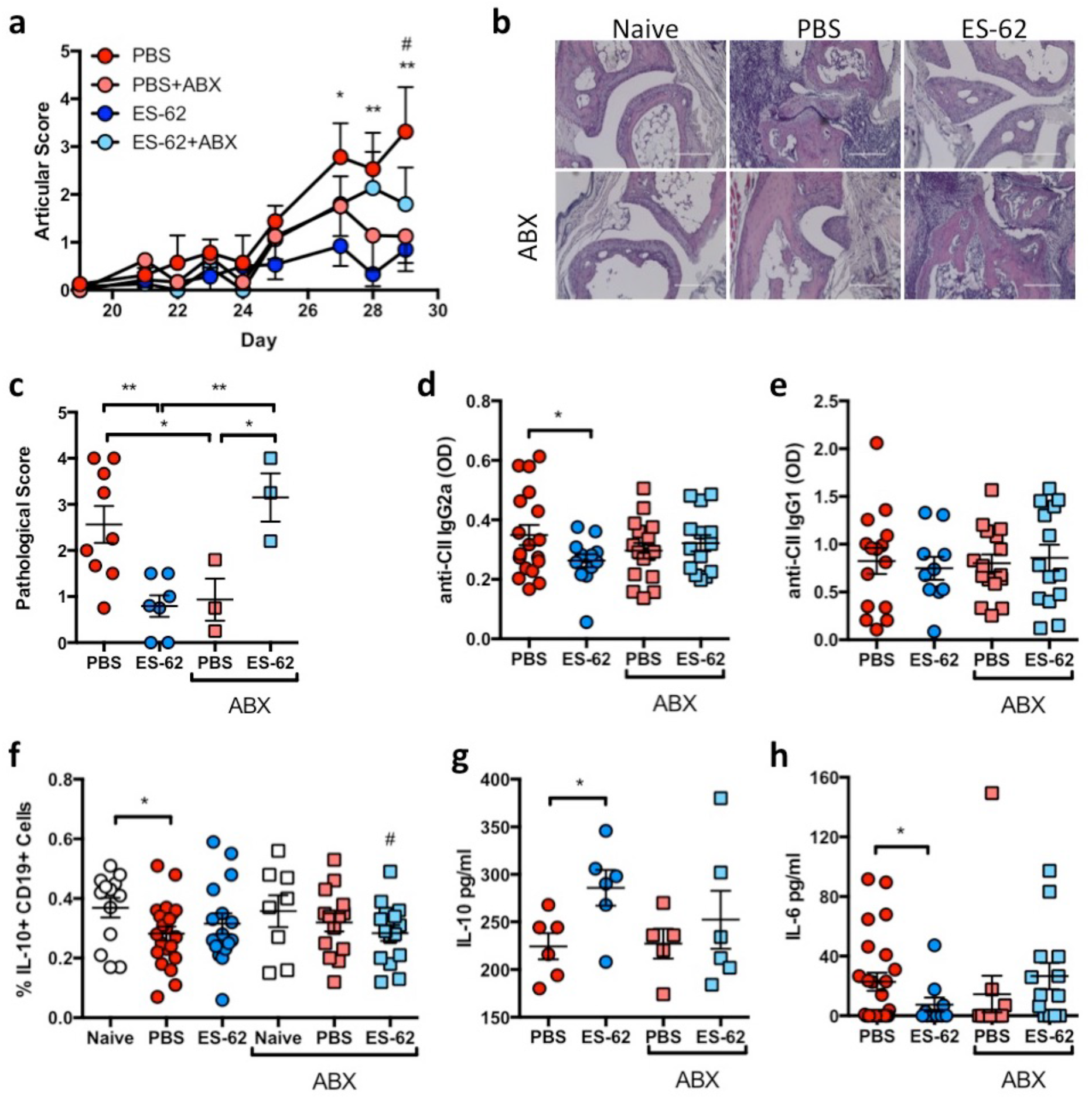
Antibiotic treatment of PBS‐ and ES-62-CIA animals results in an intermediate disease phenotype. (**a**) One week prior to CIA initiation, ABX was provided in the drinking water of DBA/1 mice and maintained throughout the course of the experiment to deplete bacteria. Data are presented as articular scores (mean ± SEM) and have been pooled from three independent experiments (PBS; n = 23, ES-62; n = 19, PBS-ABX; n = 22 and ES-62-ABX; n = 22). (**b**) H&E staining of hind paws from representative mice from each treatment group are shown on the images and scale bars represent 200 μm. (**c**) Blind scoring of the joint pathology exhibited in experimental replicates of joint sections (Clinical articular score of the individual paws were, PBS ≥ 3 (n = 9) ES-62 = 0 (n = 7), PBS-ABX = 0 (n = 3) and ES-62-ABX ≥ 3 (n = 3)). (**d** and **e**) Levels of collagen II (CII)‐ specific IgG2a (**d**; PBS; n = 18, ES-62; n = 13, PBS-ABX; n = 16 and ES-62-ABX; n = 14) and IgG1(**e**; PBS; n = 15, ES-62; n = 10, PBS-ABX; n = 16 and ES-62-ABX; n = 14) antibodies in serum were determined by ELISA. (**f**) Splenic B regulatory cells (CD19^+^IL-10^+^ cells) were determined by flow cytometry (Naïve; n = 13, PBS; n = 21, ES-62; n = 14, Naïve-ABX; n = 8 PBS-ABX; n = 14 and ES-62-ABX; n = 15). (**g** and **h**) IL-10 (**g** ‐ PBS; n = 6, ES-62; n = 5, PBS-ABX; n = 6 and ES-62-ABX; n = 6) and IL-6 (**h** ‐ PBS; n = 24, ES-62; n = 10, PBS-ABX; n = 12 and ES-62-ABX; n = 13) concentrations in serum were determined by ELISA. Statistics: all results are presented as mean ± SEM and each symbol represents an individual mouse; data are from one (**g**) or pooled from two or three independent experiments. Statistical significance was determined using two-way ANOVA (**a**), oneway ANOVA (**c** - **h**) with LSD Fishers multiple comparisons and significance indicated by asterisks, * = p < 0.05 and ** = p < 0.01.

ES-62-mediated protection against CIA is associated with restoration of the homeostatic balance of regulatory:effector B cell responses via down-regulation of aberrant MyD88 signalling^2,44^. Typically, ES-62 acts to reduce pathogenic anti-CII IgG2a but not IgG1 antibody production^45,62^. Reflecting the ABX-driven amelioration of CIA pathology, it was noted that anti-CII Ig2a levels in PBS-ABX mice are not significantly different from the lower levels of these antibodies pertaining in ES-62-CIA mice whilst those in ES-62-ABX mice are no longer significantly reduced relative to those in PBS-CIA mice (Fig 2d). No differences were detected amongst any of the group in terms of anti-CII IgG1 antibodies (Fig. 2e). Consistent with the effects of ABX on effector B cell responses, analysis of splenic IL-10+CD19^+^ “regulatory” B cells showed that both the decrease in these cells occurring during CIA (PBS-CIA) and also the maintenance of healthy levels in ES-62-CIA mice^2,44^ were lost in ABX-treated animals, with the levels in ES-62-ABX mice mirroring those reduced levels found in PBS-CIA animals (Fig. 2f). Moreover, whilst ES-62 acts to increase serum IL-10 levels in CIA mice, this regulatory cytokine is found at similarly low levels in PBS-CIA, PBS-ABX and ES-62-ABX mice (Fig. 2g). At the same time, ES-62-mediated suppression of serum levels of IL-6, a cytokine that promotes B cell (auto)immunity^63,64^, is lost following ABX treatment such that ES-62-ABX mice display the high levels of this cytokine found in PBS-CIA mice (Fig. 2h).

In addition to maintaining immune system homeostatic regulation and promoting resolution of inflammation that becomes dysregulated in CIA mice, ES-62 also acts to suppress the functional maturation of osteoclasts (OC)^43^ that directly cause erosive joint damage. Interestingly, changes in the intestinal microbiome have been shown to impact on bone mass^65-67^ and we therefore investigated whether the intermediate phenotype of joint pathology occurring in ABX-treated CIA mice reflected modulation of osteoclastogenesis resulting from perturbation of the gut microbiota. As shown previously, exposure to ES-62 *in vivo* had no effect *per se* on the numbers of OC differentiated from bone marrow (BM) progenitors *ex vivo* (Fig. 3a) but rather, it blocked their fusion to large, active multinucleated cells (Fig. 3a - c) that resorb bone^43^. Consistent with its amelioration of CIA pathology, ABX administration resulted in a decrease in large multinucleated OC and a corresponding increase in the total numbers of OCs detected in BM from CIA mice (Fig. 3a - c). ES-62 rewires osteoclastogenesis by modulating the RANK/OPG bone remodelling axis^43^ and this is evidenced again here by its ability to significantly decrease expression of RANK and increase (albeit not significantly) expression of the decoy receptor, OPG in BM relative to that seen in BM from PBS-CIA mice (Fig. 3d and e). This axis is indeed targeted by ABX treatment, with the elevated RANK expression observed in PBS-CIA BM being lost following ABX treatment such that the levels in BM from both PBS-ABX and ES-62-ABX mice were not being significantly different to those detected in ES-62-CIA BM. In addition, OPG expression was essentially the same in PBS‐ and ES-62-ABX mice and similar to that in PBS-CIA BM (Fig. 3d and e): these changes would result in similar levels of RANKL-driven osteoclastogenesis consistent with the intermediate CIA phenotype observed in PBS‐ and ES-62-ABX animals. Supporting these ABX-changes in osteoclastogenesis and consequently, bone damage in CIA, PBS-ABX and ES-62-ABX animals exhibit similar grip strengths (70.8±5.4% and 58.8±6.0%, respectively) that are intermediate to those displayed by PBS-(42.3±6.2%) and ES-62-CIA (90.1±9.6%, relative to naive mice) (Fig. 1). Our data are reminiscent of recent studies showing that commensal bacteria do not impact on differentiation of monocytes to the osteoclast lineage *per se*, but rather enhance functional maturation of OCs as evidenced by their greater size (x2.5 fold) and bone resorption capabilities. Moreover, this appears to be achieved by commensal bacteria modulating the RANKL:OPG ratio to result in increased and sustained RANK signalling^68^.

**Fig. 3.**
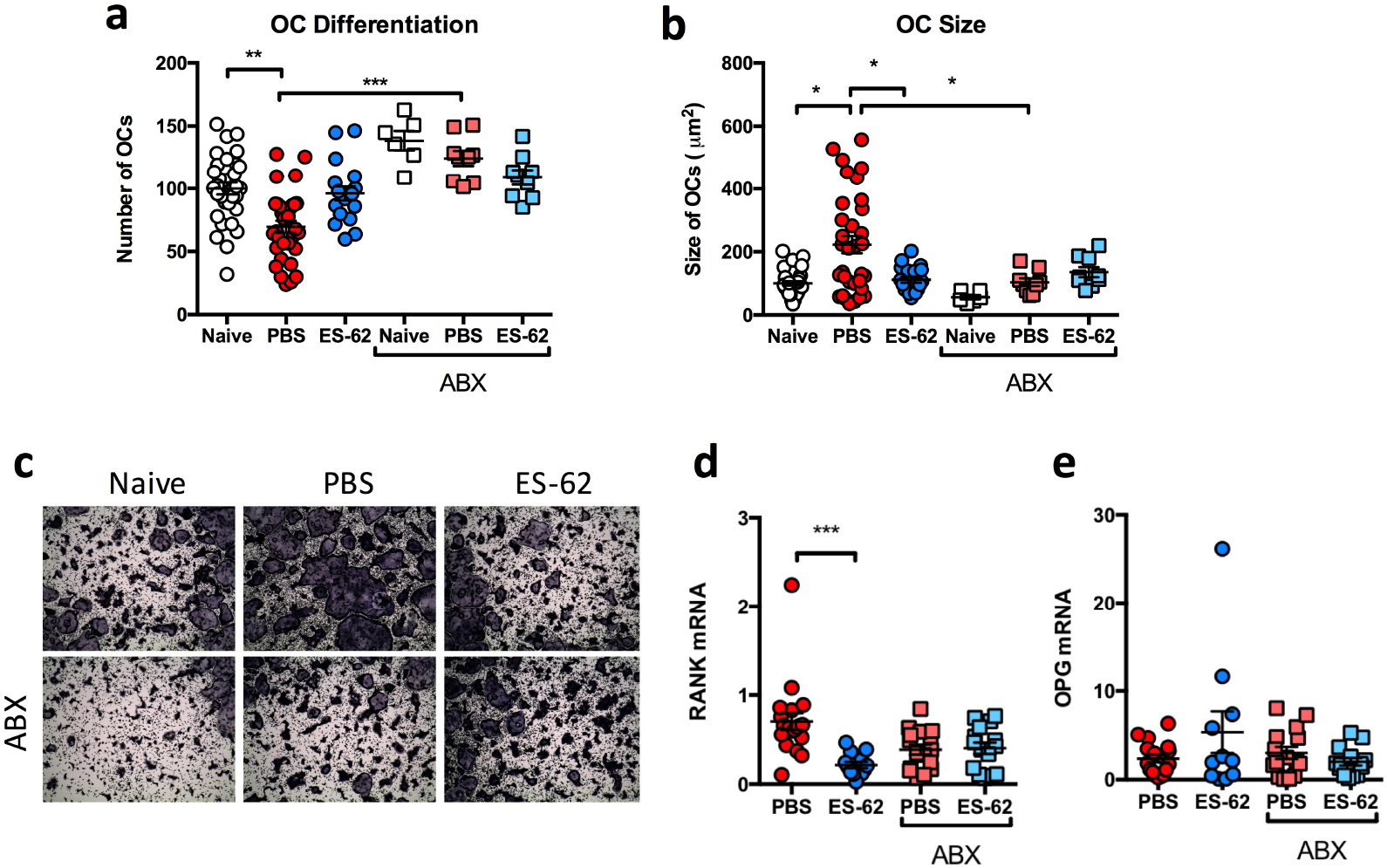
ES-62 requires the gut microbiome to protect the bone remodelling axis. (**a**, **b** and **c**) Osteoclasts were differentiated from bone marrow obtained at cull and cultured for 5 days before differentiation (**a**) and size (**b**) of osteoclasts was measured using ImageJ analysis software and data was normalised as a percentage of Naïve controls with representative images (**c**) provided (Naïve; n = 11, PBS; n = 11, ES-62; n = 6, Naïve+ABX; n = 2, PBS+ABX; n = 3, ES-62+ABX; n = 3). Whole bone marrow was used to quantify RANK (**d**; PBS; n = 20, ES-62; n = 13, PBS+ABX; n = 15, ES-62+ABX; n = 15) and OPG (**e**; PBS; n = 17, ES-62; n = 11, PBS+ABX; n = 16, ES-62+ABX; n = 14) mRNA levels using qRT-PCR and fold change was calculated following normalisation to Naïve controls. Statistics: all data are presented as mean ± SEM. In **a** and **b**, each symbol represents experimental replicates and in **d** and **e** each symbol represents individual mice and data are pooled from three independent experiments. Statistical significance was determined using one-way ANOVA (**a** and **b**) with LSD Fishers multiple comparisons or Kruskal-Wallis with Dunn’s multiple comparisons tests (**d** and **e**) and significance indicated by asterisks, * = p < 0.05, ** = p < 0.01 and *** = p < 0.001.

### 3.3. ES-62 protects against gut pathology in CIA

Intestinal dysbiosis has been associated with loss of gut integrity and chronic inflammation in conditions such as obesity that are known to promote autoimmune disorders like RA and cardiovascular comorbidities^69,70^. Our metagenomic analysis showed that established CIA was associated with changes to the gut microbiome, particularly with respect to a reduction in butyrate-producing species (Fig. 1g, i, j and S1) that have been implicated in the maintenance of epithelial integrity and gut health^50,58^. Strikingly, we have found that mice with CIA display severe gut pathology and inflammation, the levels of which directly correlate with severity of CIA (Fig. 4a). Moreover, treatment with ES-62 protects against such gut damage, specifically the thickening and generation of stubby villi in the ileum (Fig. 4b and c) and hole-like “lesions” appearing in the colon (Fig. 4b and d). The physiological relevance of these colon “lesions” is unclear but may reflect the widely-established suppression of mucus production by goblet cells in response to bacteria^71,72^ or alternatively, they may be attachment/effacement lesions induced by pathogenic bacteria including *E. coli*^73^. The gut pathology (both the ileum villi thickness and colon lesions) observed in PBS-CIA mice was abrogated in such animals administered ABX (Fig. 4c and d) suggesting that the associated reduction in chronic gut (and consequently systemic) inflammation contributes to the amelioration of CIA in PBS-ABX mice. Of note, ES-62-ABX mice displayed increased pathology and inflammatory cell infiltration of the gut tissue compared to ES-62-CIA mice, but again this was also consistent with chronic gut inflammation promoting disease severity.

**Fig. 4.**
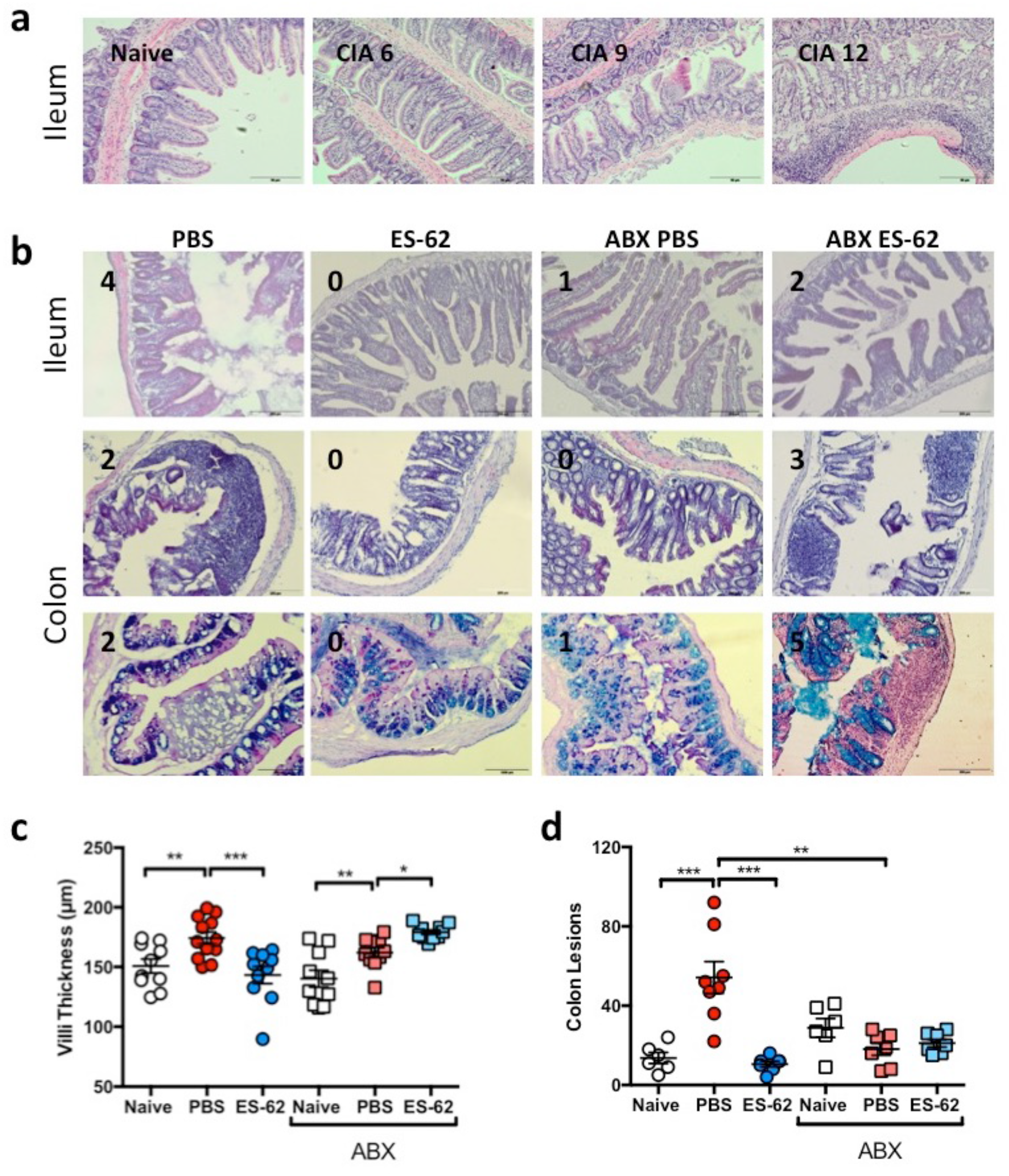
CIA is accompanied by microbiome-dependent gut pathology. (**a**) Representative H&E images (scale bar representing 50 μm) of ileum sections of naive and PBS-CIA mice with differential disease severity at cull (mouse articular score as indicated). (**b**) Representative H&E images of ileum (top row), colon (middle row) and PAS-stained colon (bottom row) sections displaying scale bar (200 μm) and articular scores of the mice are shown on the images. (**c**) Quantitative analysis of ileum villi thickness where symbols represent mean values of replicate sections as measured using ImageJ analysis software (n = 3/group with 3 – 5 replicates/animal and data are representative of two independent experiments). (**d**) Lesions were enumerated per colon section of individual mice using ImageJ imaging software (Naïve; n = 6, PBS; n = 8, ES-62; n = 7, Naive+ABX; n = 6, PBS+ABX; n = 7, ES-62+ABX; n = 7). Statistics: data are pooled from two independent experiments. Statistical significance was determined using one-way ANOVA with LSD Fishers multiple comparisons and indicated by asterisks, * = p < 0.05, ** = p < 0.01 and *** = p < 0.001.

To further address the role of gut pathology in CIA, we analysed ileum and colon tissue from naive mice and those undergoing CIA (treated with PBS or ES-62) at key points during initiation and progression of disease: (i) Naive mice, (ii) the breaking of tolerance and initiation of disease, following immunisation at day 0 with CII/CFA (≤ day 14), (iii) preclinical (day 21, prior to booster immunisation with CII) and (iv) established disease (≥ day 28; articular score: PBS – 5.2±0.8; ES-62 ‐ 0.9±1.24) phases. This analysis revealed the presence of gut pathology in PBS-CIA mice during the initiation phase with an increase in ileum villi thickness and a significant increase in the number of colonic lesions following immunisation (Fig. 5a - c). Interestingly, the ileum pathology occurring during the initiation stage in PBS-CIA mice appeared to resolve in these animals by the end of the pre-clinical phase (day 21), although thickening and shortening of the villi was again induced by the booster CII immunisation. This pattern was not the case for the colon lesions, which peaked at day 21 and were maintained throughout active disease in PBS-CIA mice. As opposed to CIA-mice, ileum integrity was maintained throughout all phases of disease in ES-62-CIA-mice whilst the high levels of colon lesions generated by day 21 of the pre-clinical phase were reduced in ES-62-treated animals (Fig. 5b and c).

**Fig. 5.**
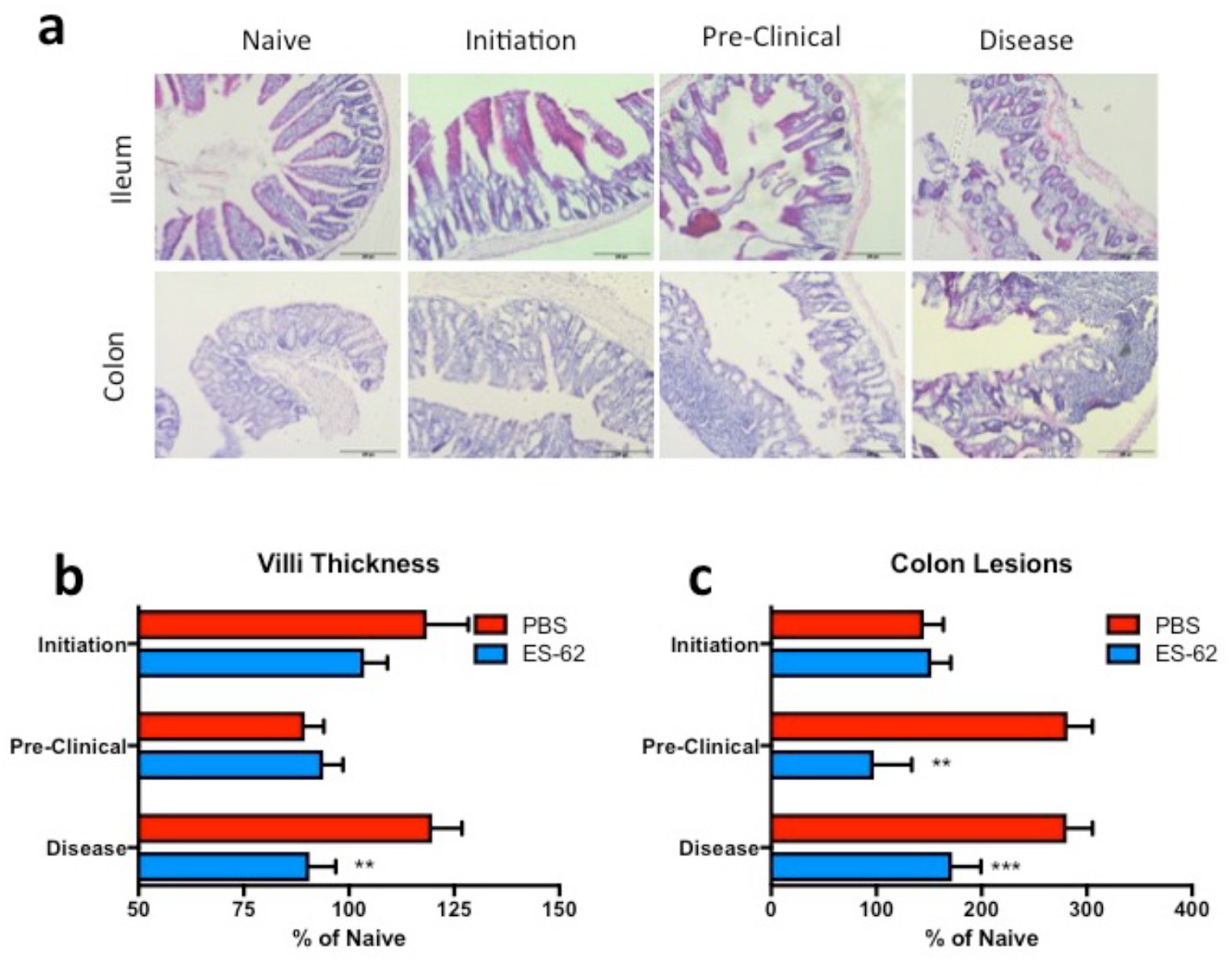
ES-62 protects against the gut pathology occurring prior to onset of arthritis. (**a**) Representative H&E images of ileum and colon pathology of mice culled during the following phases of CIA: Naïve, initiation (post-immunisation ≤d14), preclinical (d21 prior to challenge) and disease (established arthritis d≥28, articular score PBS – 5.2±0.8; ES-62 – 0.9±1.24). Scale bars are 200 μm. (**b** and **c**) Changes in the villus thickness (**b**; Initiation; PBS ‐ n = 5, ES-62 ‐ n = 6, Pre-Clinical; n = 3, Disease; PBS ‐ n = 4, ES-62 ‐ n = 5 mice) and number of colon lesions (**c**; Initiation; n = 6, Pre-Clinical; n = 3, Disease; PBS ‐ n = 4, ES-62 ‐ n = 5 mice) were quantified using ImageJ analysis software at each phase of the experiment and data were normalised to values of naïve mice with representative images displayed. Statistics: data are presented as mean ± SEM of individual mice from one experiment. Statistical significance was determined using two-way ANOVAs to compare PBS and ES-62 treatment at each time point (**b** - **c**) and significance is denoted as * = p < 0.05 and *** = p < 0.001.

Validating our metagenomic analysis, qPCR analysis showed that ES-62 acts to prevent the enrichment of Bacteriodetes and maintain levels of Firmicutes in the colon of mice during established CIA (Fig. 6a and b). However, analysis at the various pre-clinical phases of the disease showed a more dynamic situation: for example, following the primary immunisation with CII/CFA, there is a significant decrease in the levels of both Bacteroidetes and Firmicutes, with the rise in Bacteriodetes evident in established arthritis only occurring following the booster immunisation (Fig. 6a and b). In contrast, ES-62 acts to maintain “healthy” levels of Firmicutes in CIA-mice throughout but most strongly during the initiation and preclinical phase where it also acted to maintain the lowest levels of Bacteroidetes (Fig. 6a and b). Moreover, although ES-62 treatment promoted enrichment of butyrate-producing bacteria (*Butyrivibrio*) at all stages of disease, this was most evident in the initiation phase of disease, and presumably this promotes and maintains gut integrity (Fig. 6c).

**Fig. 6.**
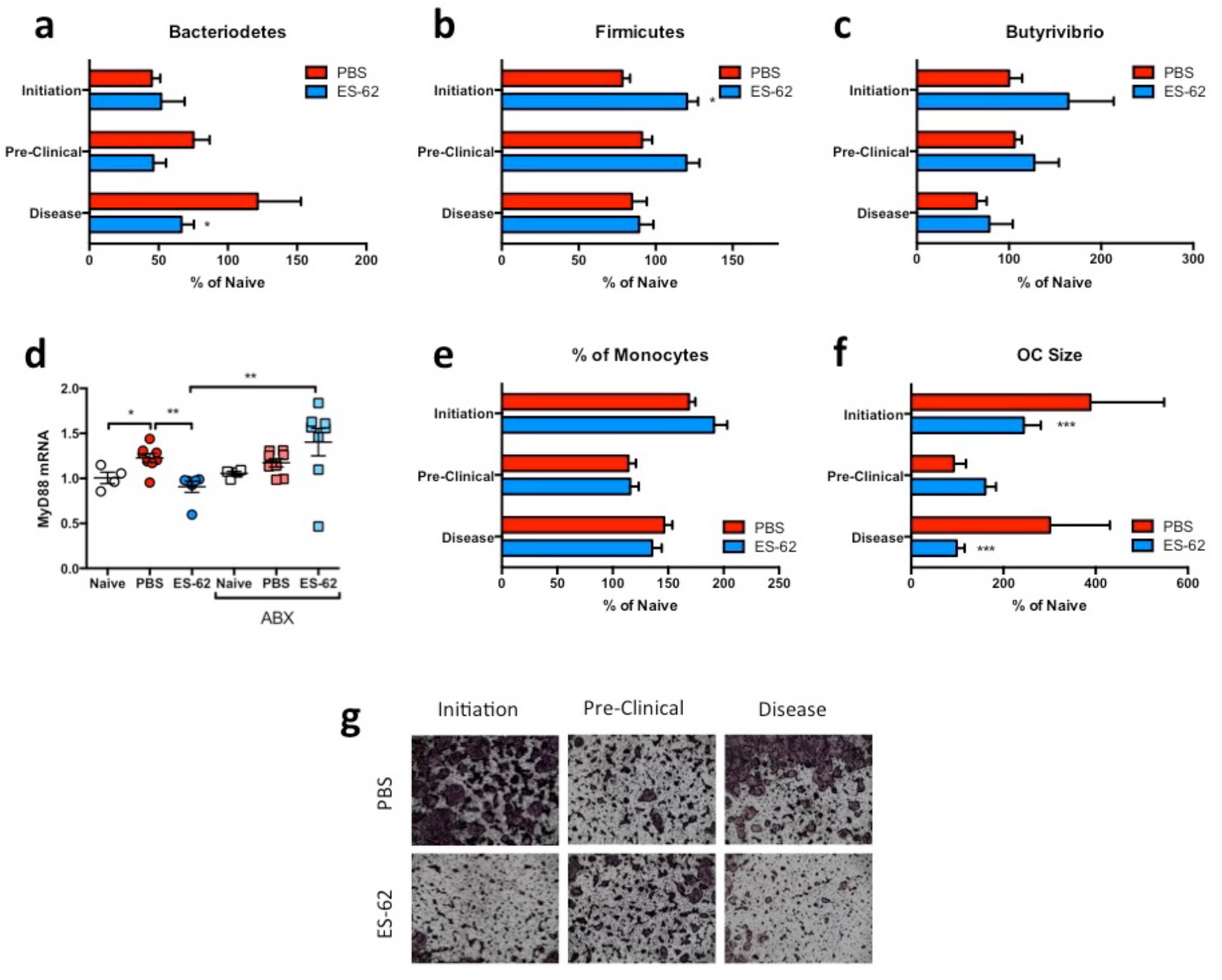
ES-62 modulates the gut-bone marrow axis during the early phases of CIA. (**a**, **b** and **c**) Changes in Bacteriodetes (**a**), Firmicutes (**b**) and Butyrivibrio (**c**) populations in the colon faecal matter of Naïve, PBS‐ or ES-62-treated animals were measured at each stage of disease by qPCR and data were normalised to total bacterial content and presented as % change compared to Naïve mice. (**d**) Whole bone marrow was used to quantify MyD88 mRNA levels using qRT-PCR and normalised to Naïve controls. (**e**) The proportion of monocytes (CD3^+^B220^−^Ter119^−^Ly6G^−^Ly6C^+^) in bone marrow was measured by flow cytometry and normalised to those in Naive control mice. (**f** and **g**) Osteoclasts were differentiated from bone marrow obtained at cull and cultured for 5 days and size of osteoclasts (**f**) was measured using ImageJ analysis software and normalised to those from naive controls with representative images for each disease stage in both treatment groups displayed (**g**). Statistics: data are presented as mean ± SEM values from individual animals (**a**, **b** and **c**; Initiation; n = 6, Pre-Clinical; n = 3, Disease; n = 10. **d**; Naïve; n = 4, PBS; n = 8, ES-62; n = 6, Naive+ABX; n = 4, PBS+ABX; n = 8, ES-62+ABX; n = 8. **e**; Initiation; n = 6, Pre-Clinical; n = 3, Disease; n = 4) or mean ± SD from experimental replicates (**f**; Initiation; n = 18, Pre-Clinical; n = 9, Disease; PBS ‐ n = 12 and ES-62 – n = 15). Statistical significance was determined using two-(**a**, **b**, **c**, **e** and **f**) or one-way (**d**) ANOVA with LSD Fishers multiple comparisons and indicated by asterisks, * = p < 0.05, ** = p < 0.01 and *** = p < 0.001.

Commensal bacteria play key roles in educating the immune system and bone marrow progenitors and hence, loss of diversity in the microbiome can impact on both inflammation and bone homeostasis^68,74^. Such “training” involves interactions of Pattern Recognition Receptors (PRR; e.g. TLRs and NODs) with the gut microbiome^2,5,13,75^. Consistent with this, ES-62 can rewire BM progenitors and stromal cells from CIA mice to an anti-inflammatory, regulatory or tissue repair phenotype^2,39,43,46,76,77^ by subverting TLR4 signalling to prevent the up-regulation of the key TLR signal transducer, MyD88 observed during chronic inflammation^78,79^. Interestingly, therefore, given the increased incidence and severity of CIA in ES-62-ABX mice, we show that ES-62 dampening of aberrant MyD88 expression is abolished by ABX treatment and indeed, that MyD88 is expressed at equivalent levels in BM from PBS-CIA, PBS-ABX and ES-62-ABX mice (Fig. 6d).

Commensal bacteria can also dynamically educate OC progenitor (OCP) maturation^68,74^: as OCs normally act in concert with osteoblasts (OBs) to homeostatically maintain healthy bone, the enhanced functional capacity of OCs elicited by commensal bacteria may actually render hosts more susceptible to bone damage during chronic inflammation, a condition that promotes bone remodelling, particularly as commensal bacteria also appear to act to suppress OB function^68,74^. Reflecting the dynamic and differential changes in colonic microbiota, gut inflammation and pathology, there is a strong increase in the monocyte populations containing OCPs during the early inflammatory initiation phase of CIA that has resolved by the end of the preclinical phase only to increase again in the established phase of arthritis following the booster CII immunisation. Consistent with our data that ES-62 does not fundamentally suppress OC differentiation, there are no differences in the percentage of monocytes between the PBS-CIA and ES-62-CIA groups (Fig. 6e). However, BM from CIA mice following both primary and booster immunisations, shows enhanced capacity for functional OC maturation that is suppressed by *in vivo* exposure to ES-62 (Fig. 6f and g).

## 4. Discussion

This study is the first to demonstrate that a defined, systemically-acting parasitic worm-derived product can impact on, and harness, the microbiome to exert its therapeutic effects against chronic inflammation in target organs distal to the gut such as the joints. Collectively, our data suggest that ES-62-mediated protection against inflammatory joint disease is associated with normalisation of the gut dysbiosis observed in arthritic mice towards the “healthy” microbiome observed in naive mice. This “normalisation” is highly reminiscent of that seen following treatment of RA patients with DMARDs^80^ and lends further weight to increasing evidence that changes in microbiome status may contribute to pathogenesis in this multifactorial disease^31,49,74^,^81,82^. Indeed, perturbation of the microbiota has been shown to influence both the generation of pathogenic Th17 cells and also the homeostatic induction of inflammation-resolving Bregs and Tregs in experimental models of RA and human disease^20,24-26^. Moreover, and consistent with the idea that microbiome status can promote disease in susceptible individuals, development of arthritis in a number of animal models is lost under “Germ-Free” (GF) conditions or following depletion of the microbiota by treatment with broad-spectrum ABX25,49,59,61.

Certainly, potentially pathogenic changes in the gut microbiome have been reported following analysis of stool faeces from patients with new-onset disease as well as those with established arthritis. However, differential, and even contradictory, associations have been reported that perhaps reflect the difficulties in standardising stool samples or their failure to fully recapitulate the contents of the gut microbiota and the specific communities shaped by the intestinal microenvironment ^31^,^83-85^. Nevertheless, with respect to the Bacteroidetes, whose levels we found to be generally elevated during established CIA, *Prevotella copri* has been reported to be enriched in stool samples of new-onset patients^24,86^. Moreover, transfer of such faecal matter containing *Prevotella copri* into GF arthritis-prone SKG mice leads to enhanced levels of zymosan-stimulated intestinal Th17 cells and more severe arthritis^86^. However, SKG mice are increasingly perceived to be a model of Spondyloarthritis rather than RA^31^ and perhaps reflecting this, although we found *Prevotellaceae* species to be upregulated in the colon of mice with established arthritis, this was neither restricted to *P. copri* nor was it reversed by treatment with ES-62. Moreover, *P. histocola* has been reported to be protective against CIA in mice transgenic for the RA-risk gene HLA-DQ8^87^, whilst at the genus level, *Prevotella* species were found to be under-represented in the IL-1rn^−/−^ model of autoimmune arthritis^60^. In terms of the pathogenic potential of Bacteroidetes in CIA, we found *B. vulgatus* to be significantly enriched and *A. putredinis* (*Rikenellaceae* family) to be depleted in the colon of CIA relative to Naive or ES-62-CIA mice. Interestingly, a similar inverse pattern of *B. vulgatus* and *Rikenellaceae* species was observed in rats transgenic for HLA-B27, a humanised rodent model for rheumatological conditions such as Ankylosing Spondylitis, Reactive Arthritis and Psoriatic Arthritis^88^.

Colonisation with segmented filamentous bacteria (SFB) from the Firmicutes phylum has likewise previously been shown to promote (IL-22-dependent^89^) Th17 responses and arthritis in GF K/BxN mice^20^. However, as also reported for the IL1rn^−/−^ model^60^, we were unable to detect any OTUs that could be identified as SFB (e.g. *Candidatus arthromitus*) in our metagenomic analysis: moreover, we detected no significant differences by qPCR in the levels of SFB found in either the colon and ileum between PBS‐ and ES-62-treated CIA mice at any phase of disease (data not shown). Our findings therefore resonate with the report that despite being present in all cages, SFB could only be detected in 2/10 mice undergoing CIA prior to the booster CII immunisation^61^ and of relevance to RA, does not appear to be present in the gut metagenomic analysis of the large numbers of adult human faecal samples sequenced for the Human Microbiome Project^85^. Indeed, and further questioning the role for SFB in the K/BxN model, a recent study reports that IL-17 actually appears dispensable for arthritis in K/BxN mice and proposes that the gut microbiota regulates joint disease via impacting on T follicular helper (Tfh) rather than Th17 cells^90^.

Further faecal profiling of Firmicutes has linked changes in the *Clostridiaceae*, *Coriobacteriaceae* and *Lachnospiraceae* families to patients with established disease^80,82^. Our metagenomic analysis of luminal colon and ileum faecal matter shows the protective actions of ES-62 to be most strongly associated with maintenance of the *Clostridiaceae*, *Lachnospiraceae and Ruminococcaceae* families, particularly those associated with butyrate production (*Blautia*, *Roseburia*, *Dorea* and *Butyrvibrio* and *Ruminococcus*). Interestingly, *Ruminococcus* species were found to be depleted in the IL1rn^−/−^model of inflammatory arthritis supporting the suggestion that maintenance and/or enrichment of butyrate-producing bacteria are protective^91^. Reflecting this, administration of butyrate was found to ameliorate severity of CIA, notably in terms of reduced inflammatory cell infiltration of the joint, pannus formation and cartilage and bone destruction^91^. In contrast, administration of butyrate exacerbated antibody-induced arthritis, a model wherein the administration of serum from K/BxN mice (containing anti-glucose-6-phosphate isomerase autoantibodies) to C57BL/6 mice bypasses the initiation and adaptive immunity phases of disease. These apparently conflicting data can be reconciled by the proposal that butyrate needs to act during the preclinical phases of disease to exhibit its protective actions^91^. Perhaps consistent with this, whilst ES-62 protects against the depletion of butyrate-producing species observed in mice with established disease, we also find *Butyrivibrio* to be most enriched by ES-62 in the initiation phase of CIA. Collectively, these data suggest that depletion of butyrate-producing bacteria associated with the onset of CIA may contribute to the gut pathology promoting and perpetuating the inflammation that drives the breakdown of immune tolerance and consequent autoimmune joint disease.

Certainly, we find gut pathology to precede onset of joint disease, being detectable within 6 days of the primary CII immunisation and the loss of colon barrier integrity peaking by the end of the preclinical phase (d21) of CIA. Moreover, such gut pathology is accompanied by dynamic changes in the microbiome of CIA-mice as evidenced by the decrease in the colon abundance of Bacteroidetes and Firmicutes during the early initiation phase of disease and the enrichment of the former in mice during established arthritis. A similar depletion of Bacteriodetes during the preclinical phase (d21) of CIA has been independently reported but in this case, was accompanied by a corresponding enrichment of Firmicutes^61^. Our failure to observe such an enrichment of Firmicutes at this time point may simply reflect our analysis of colon contents rather than faecal matter. Rather, we found ES-62 maintained and enhanced the colon levels of Firmicutes throughout all phases of CIA, likely by promoting enrichment of Butyrate-producing *Lachnospiraceae* species as seen in our metagenomic analysis of both ileum and colon faecal matter in mice with established arthritis. In any case, the dynamic and intestinal-site selective nature of the dysbiosis observed in CIA, and the protection against it afforded by ES-62, underlines the need to identify and validate precise protective microbial signatures and the context of their complex biogeographic microenvironment^84^, for effective therapeutic intervention at all stages of human RA.

Butyrate is known to regulate gut barrier integrity^92,93^ and goblet cell production of MUC2^94^ and hence depletion of butyrate-producing species during CIA likely promotes and perpetuates dysbiosis and loss of gut barrier integrity. One potential consequence of this gut pathology is the aberrant systemic colonisation of pathogenic bacteria as illustrated here in CIA by the outgrowth of *Proteobacteria* [particularly *E. coli* and *H. hepaticus*, the latter of which has also been reported to be enriched in the IL1rn^−/−^ model^60^] and accompanying gut pathology reminiscent of the suppression of goblet cell mucus production and attachment/effacement lesions resulting from pathological enteric infections, including *E. coli* ^71-73^. Perhaps reflecting the loss of diversity in the microbiota and the breakdown of intestinal homeostasis in RA, in addition to their increased faecal abundance of *P. copri*, early onset RA patients also exhibit *P. copri* in their synovial fluid^24,95^. Moreover, *E. coli* and other infectious microorganisms have been reported to commonly colonise RA patients and may exacerbate disease^96-98^.

The normalisation of the gut microbiota by ES-62 may actually result in a dual pronged mechanism by which butyrate, in addition to its local gut-protecting actions, could also impact systemically to protect more directly against joint pathology in CIA. Consistent with this idea, butyrate has been reported to suppress osteoclastogenesis^99^ and by protecting against pathological bone loss, to regulate bone mass^65^. Intriguingly, we also find spikes of functional maturation of OCs in the initiation and established phases of CIA that are prevented by ES-62 and are associated with its enrichment of butyrate-producing species in CIA-mice. The mechanisms by which ES-62 orchestrates such maintenance of the complex homeostasis of the gut-bone marrow axis are not clear. However, it is intriguing given that TLR4/MyD88 signalling is the primary target of ES-62 in promoting Bregs and suppressing Th17-mediated inflammation^2,39,41^,^44^, that the systemic Th17 differentiation and consequent autoimmune arthritis occurring in the IL1rn^−/−^ model is dependent on TLR4^100^. Moreover, the accompanying dysbiosis, that on faecal transfer can confer arthritis-predisposing Th17 inflammation in wild type mice, is also regulated by TLR4^60^. Furthermore, interestingly, 11/44 taxa disrupted in IL1rn^−/−^ mice were normalised in IL1rn^−/−^TLR4^−/−^ animals and these included *Ruminococcus* species, which are also promoted by ES-62^60^. In mechanistic terms, the resetting of Bregs levels by ES-62 in CIA has also been reported to be regulated by the microbiome in other models of autoimmune arthritis^25^. Indeed, consistent with the proposal that Bregs are homeostatically induced to resolve dysbiosis-induced inflammation in autoimmune arthritis, as indicated by disruption of the process by ABX treatment^25^, we have similarly found the ES-62-mediated restoration of IL-10^+^ B cells in CIA to be compromised by such perturbation of the microbiome. Collectively, these findings suggest that ES-62 may achieve both its immunoregulatory and microbiome normalisation effects by targeting MyD88.

TLR4/MyD88 signalling in RA had previously been attributed solely to recognition of DAMPs in the joint^78,79^ and thus collectively, these findings shed new light on its pathogenic roles in initiation and progression of disease as well as emphasise its potential as a therapeutic target in RA. In particular, they underscore the complex and central role of TLR4/MyD88 signalling in regulating the gut-bone marrow axis in musculoskeletal homeostasis and it’s dysregulation resulting in systemic inflammation, breaking of tolerance, aberrant osteoclastogenesis and consequently joint destruction in arthritis. Moreover, they suggest that ES-62 may achieve its protective effects in CIA by directly targeting this key regulatory node in order to rebalance the gut-bone marrow axis and limit aberrant inflammation and joint damage, by homoestatically restoring levels of Bregs and resetting osteoclastogenesis. Thus, by exploiting ES-62 as a unique tool to dissect pathogenic and protective microbial signatures in CIA we could potentially understand how to elicit homeostatic regulation of gut and resolve inflammation in autoimmune inflammatory arthritis.

## 5. Acknowledgements

The work was funded by an award to MMH, WH and PAH from Arthritis Research UK (21133).

## 6. Author contributions

JD, AT, MAP, FL, JC and AMK performed the experiments for the study designed by MMH, WH and PAH. JD and FL manufactured ES-62. MMH, WH and JD wrote the paper and all authors were involved in reviewing and revising the manuscript and have approved the final version.

---

**Supplementary Fig. 1.**
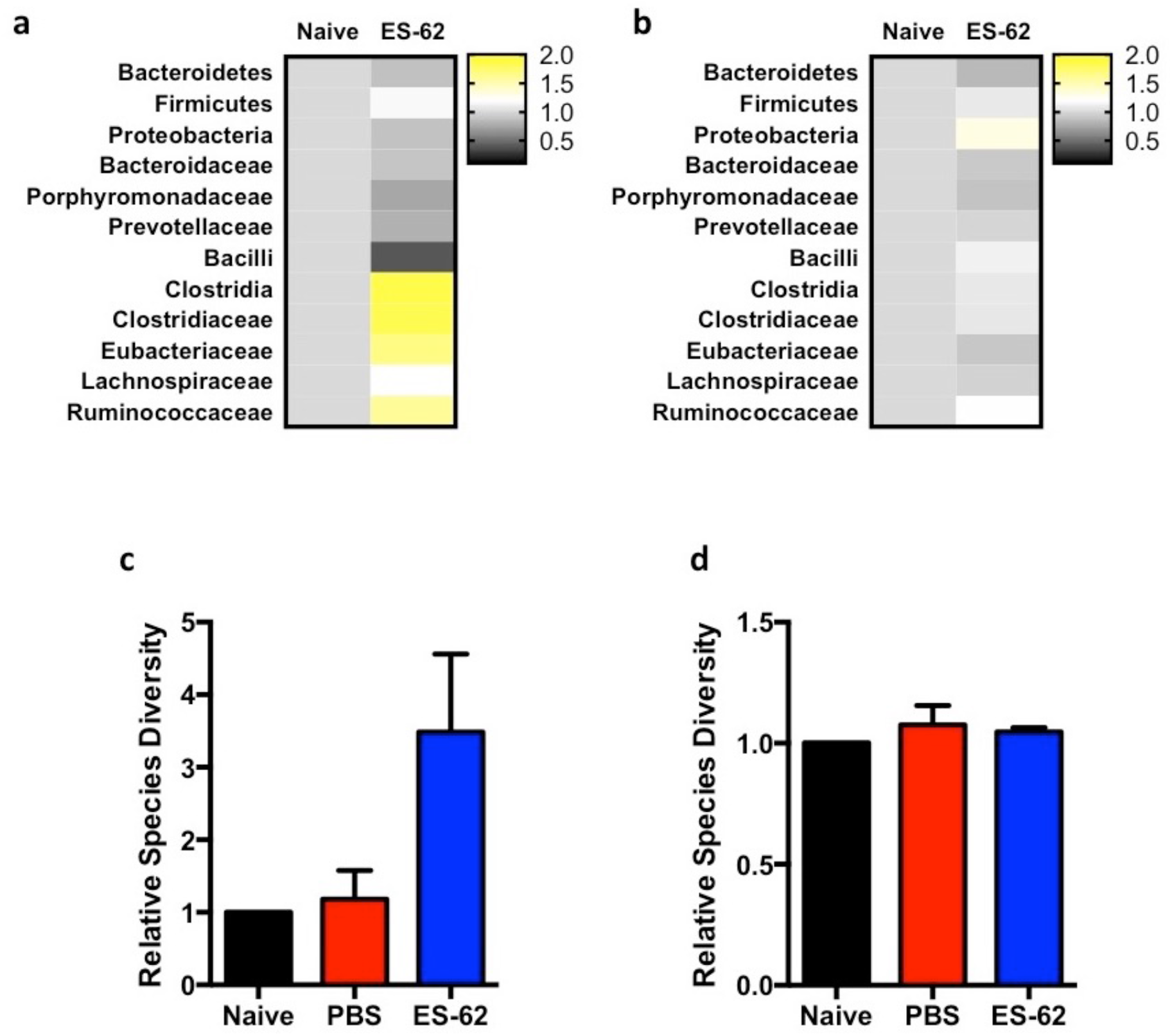
Treatment of naïve mice with ES-62 increases butyrate-producing bacteria in the ileum, but not the colon of animals. (**a** and **b**) Ileum (**a**) and colon (**b**) fecal matter from Naïve or ES-62-treated Naïve animals was analysed for changes in bacterial populations in a single experiment using samples pooled from three animals per group and displayed as Heatmaps, where the ES-62-treated samples are normalised to the Naïve controls. (**c** and **d**) The number of species detected during metagenomic analysis of the ileum (**c**) and colon (**d**) of Naive, PBS-CIA and ES-62-CIA mice. Statistics: data are presented as mean ± SEM where for metagenomic analysis, three mice per group were pooled from three independent experiments and normalised to Naïve controls (**c** and **d**).

**Supplementary Fig. 2.**
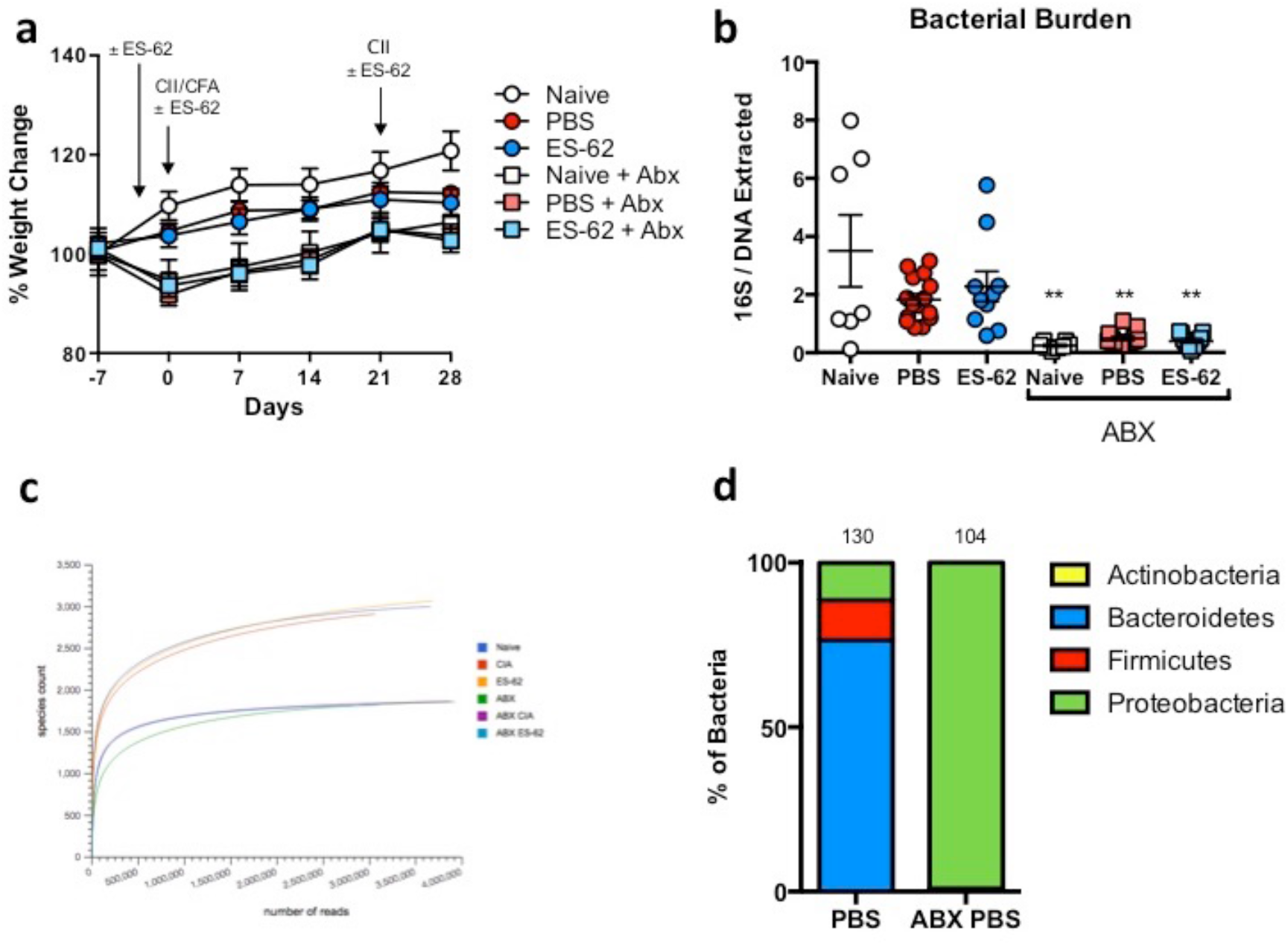
ABX treatment eliminates commensal bacteria and enriches residual proteobacteria species. (**a**) Body weights of individual animals over the time course of CIA were measured from day ‐7 (prior to ABX administration) and presented as mean % weight ± SEM (Naïve; n = 8, PBS; n = 16, ES-62; n = 13, Naive+ABX; n = 8, PBS+ABX; n = 16, ES-62+ABX; n = 15). (**b**) DNA was isolated from the colon contents of individual animals and the levels of 16S bacterial DNA detected by qPCR was measured and normalised to the levels of total DNA extracted. Symbols represent values from individual mice from a single experiment. (**c**) Representative rarefaction plots from the MG-RAST metagenomic analysis showing the levels of bacterial diversity of colon faecal matter in each treatment group from one of the three experiments. (**d**) Composition of bacterial phyla present in PBS or PBS-ABX animals (metagenomic data obtained by pooling samples from 3 mice/group from a single experiment) represented as a % of all bacteria detected. Statistics: data are presented as mean or mean ± SEM (**a**, **b** and **d**). Statistical significance was determined using one-way ANOVA with Dunn’s multiple comparisons comparing Naïve, PBS or ES-62-treated groups to their respective ABX-treated control and indicated by asterisks ** = p < 0.01.

